# Localized control of protein phase separation via membrane binding

**DOI:** 10.1101/2025.11.14.688424

**Authors:** Isabel LuValle-Burke, Giacomo Bartolucci, Daxiao Sun, Xueping Zhao, Christoph A. Weber, Alf Honigmann

**Affiliations:** Max Planck Institute of Molecular Cell Biology and Genetics, Dresden, Germany; Department of Condensed Matter Physics, Universitat de Barcelona, 08007 Barcelona, Spain; Universitat de Barcelona Institute of Complex Systems (UBICS), Universitat de Barcelona, 08028 Barcelona, Spain; Technische Universität Dresden, Biotechnologisches Zentrum, Center for Molecular and Cellular Bioengineering (CMCB), Dresden, Germany; School of Mathematical Sciences, University of Nottingham, Ningbo, China; Faculty of Mathematics, Natural Sciences, and Engineering: Institute of Physics, University of Augsburg, Germany; Cluster of Excellence Physics of Life, TU Dresden, Dresden, Germany

**Author notes:** I. L.-B. and G. B contributed equally to this work.

## Abstract

Phase separation of protein-rich condensates is key for the spatial organization of cells. Multivalent proteins can phase separate in the bulk cytoplasm to form 3D condensates or at the membrane surface to form 2D condensates. How cells control 2D versus 3D phase transitions is not well understood. Here, we combine an *in vitro* model system of membrane surface phase separation with thermodynamic modelling to explore the relation of 2D and 3D phase transitions. We engineered the phase-separating protein FUS to bind to membranes and quantified 2D and 3D phase separation as a function of total FUS concentration and salt concentration. We find that membrane binding induces the formation of 2D surface condensates far below the bulk saturation concentration. In addition to direct FUS-membrane binding, surface condensates sequester more FUS molecules from the bulk via protein-protein interactions, resulting in a substantial deviation from the classical binding isotherm. By changing the protein-protein interaction strength via salt titration, we find that the saturation concentrations for 2D and 3D phase separation are coupled. By extending the classical theory of condensate wetting via membrane binding, we recapitulate the experimental results and show that tuning the membrane-binding strength of a phase-separating protein provides a robust way to control 2D condensation at the membrane over a wide concentration range without entering the 3D condensation regime. Taken together, our results provide a simple framework to understand how cells tune protein interactions and membrane binding to control 2D and 3D phase transitions.

## Introduction

Cells must spatially organize their contents to ensure distinct biochemical environments. They can accomplish this in two ways: the implementation of lipid bilayers that act as a physical barrier and form membrane-bound organelles and vesicles, or through the formation of membrane-less compartments with defined molecular compositions, via the process of phase separation^1–4^. Over the past decade, many membrane-less compartments, including the nucleolus, stress granules, and synaptic densities, have been shown to form through phase separation and have been termed biomolecular condensates^3^. The molecular driving force behind the phase separation of biological macromolecules is multivalency, which can arise through RNA binding, intrinsically disordered regions (IDRs), or folded interaction domains^5,6^. These multivalent interactions drive the assembly of monomers into larger, more complex oligomeric species that reduce solubility and promote phase separation^3,7^. The diversity of multivalent and intrinsically disordered proteins within the proteome, combined with the rising interest in phase separation, has led to the discovery of many proteins that can undergo phase separation.

Cellular compartments formed via phase separation are often dynamically assembled and dissolved to perform their biological functions. Examples are stress granules^8–10^, P-granules^11–13^ and membrane signaling hubs^14–17^. To achieve these transitions, cells must find ways to cross the boundary between the phase coexistence region and the homogeneous region of the phase diagram in a robust and controlled manner^18–20^. The key parameters that control such phase transitions in biological systems are the strength of interactions and the concentration of the phase-separating protein(s). Both parameters are tuned by cells to control phase separation. For example, interaction strength can be modulated by post-translational modifications^21^, pH changes, ionic strength, and others, while protein concentration can be adjusted by balancing protein production and degradation^22^. A special case is when phase separation happens at the membrane surface. Membranes are, by nature, complex biochemical hubs because of their heterogeneous composition of different lipid species^23^. On membranes, the phase transitions are often triggered by recruiting a multivalent protein from the cytoplasm to the membrane, for example, via binding to activated receptor proteins^24^. Membrane binding increases the local protein concentration and limits the diffusion of molecules to two dimensions, thereby facilitating molecular interactions and leading to surface phase separation at lower concentrations than in the bulk^25^.

The interplay between membranes and membrane-less compartments formed via phase separation has recently received increased attention^26–28^, particularly in the context of reconstituting signaling and adhesion complexes on synthetic lipid bilayers^29^. Prominent examples have been reconstitutions of linker for activation of T Cells (LAT)^14,30^, the nephrin-NcK-WASP system^31,32^, and the tight junction scaffold ZO1^33^. Concepts of prewetting and surface phase separation have been used both experimentally^33–38^ and theoretically^39–41^ to assess the role of membrane binding in the facilitation of phase separation in sub-saturating bulk concentrations. However, how 2D surface condensation and 3D bulk condensation of a multivalent protein are quantitatively related remains unclear.

In this paper, we use a minimal *in vitro* system containing the multivalent protein Fused in Sarcoma (FUS) that we designed to bind to a model membrane to investigate the relationship between 2D and 3D phase separation. We find that membrane binding induces 2D condensates to emerge at a protein concentration far below the bulk saturation concentration of FUS *c*_sat_ (Figure 1). Above *c*_sat_, surprisingly, partially wetting droplets coexist with 2D patterns, giving rise to three phases that coexist stably on the membrane surface. By tuning the strength of protein interactions via salt titration, we find that the saturation concentrations for 2D and 3D phase separation are coupled. By extending the classical theory of condensate wetting by membrane binding, we recapitulate the experimental results and show that tuning the membrane binding strength of a phase-separating protein provides a robust way to control 2D condensation at the membrane over a wide concentration range without entering the 3D condensation regime. In summary, these results suggest an effective way for cells to regulate 2D and 3D phase transitions by tuning protein interactions and membrane binding.

**Figure 1.**
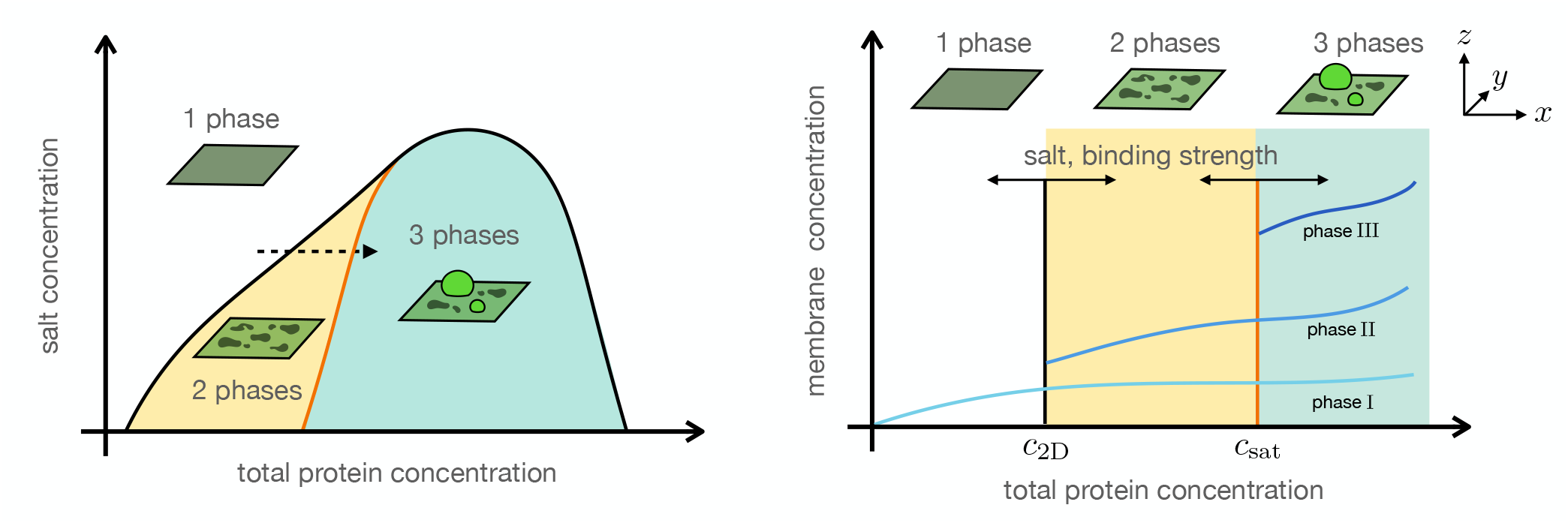
Phase separation coupled to membrane binding leads to surface patterns and three-phase coexistence. When a phase-separating protein can bind to a membrane, surface phase separation can occur for surface concentrations above the surface saturation concentration *c*_2D_ [units: molecule number per surface area]. Surface phase separation can occur for bulk concentrations far below the bulk saturation concentration *c*_sat_ [units: molecule number per volume] since molecules can bind and thus enhance the local concentration at the membrane surface. For bulk concentrations above *c*_sat_, three phases can stably coexist at the membrane surface. The respective saturation concentrations *c*_2D_ and *c*_sat_ can be tuned via salt concentration and binding strength.

## Results

### Membrane binding induces 2D phase separation in sub-saturating conditions

To elucidate the interplay between binding to a biological membrane and phase separation, we employed the well-characterized protein Fused in Sarcoma (FUS) as our model protein. FUS has been found to undergo homotypic phase separation with a saturation concentration in low micro-molar range in the absence of crowding agents *in vitro*. In addition, due to previously established understanding of the molecular grammar that governs phase separation of FUS *in vitro*, mutants are available that abrogate phase separation, such as the YtoS mutant^42^. We therefore, generated two constructs expressing the WT and the YtoS mutant linked to a GFP for visualization and a His10 tag for specific binding to the receptor-like nickel-chelated lipid DGS-NTA (Figure 2a-b). To assess the effects of the additional tags on the characterized^42^ propensity of the proteins to undergo phase separation, a bulk phase separation assay was performed on the purified proteins (Figure 2c-e) under physiological conditions (150 mM KCl pH 7.25). By assessing the fraction of condensed protein, we were able to ascertain that the two constructs’ behaviours remained in agreement with previously published results^42^. The WT construct had a bulk saturation concentration *c*_sat_ of approximately 2µM, above which condensates formed in the bulk.

**Figure 2.**
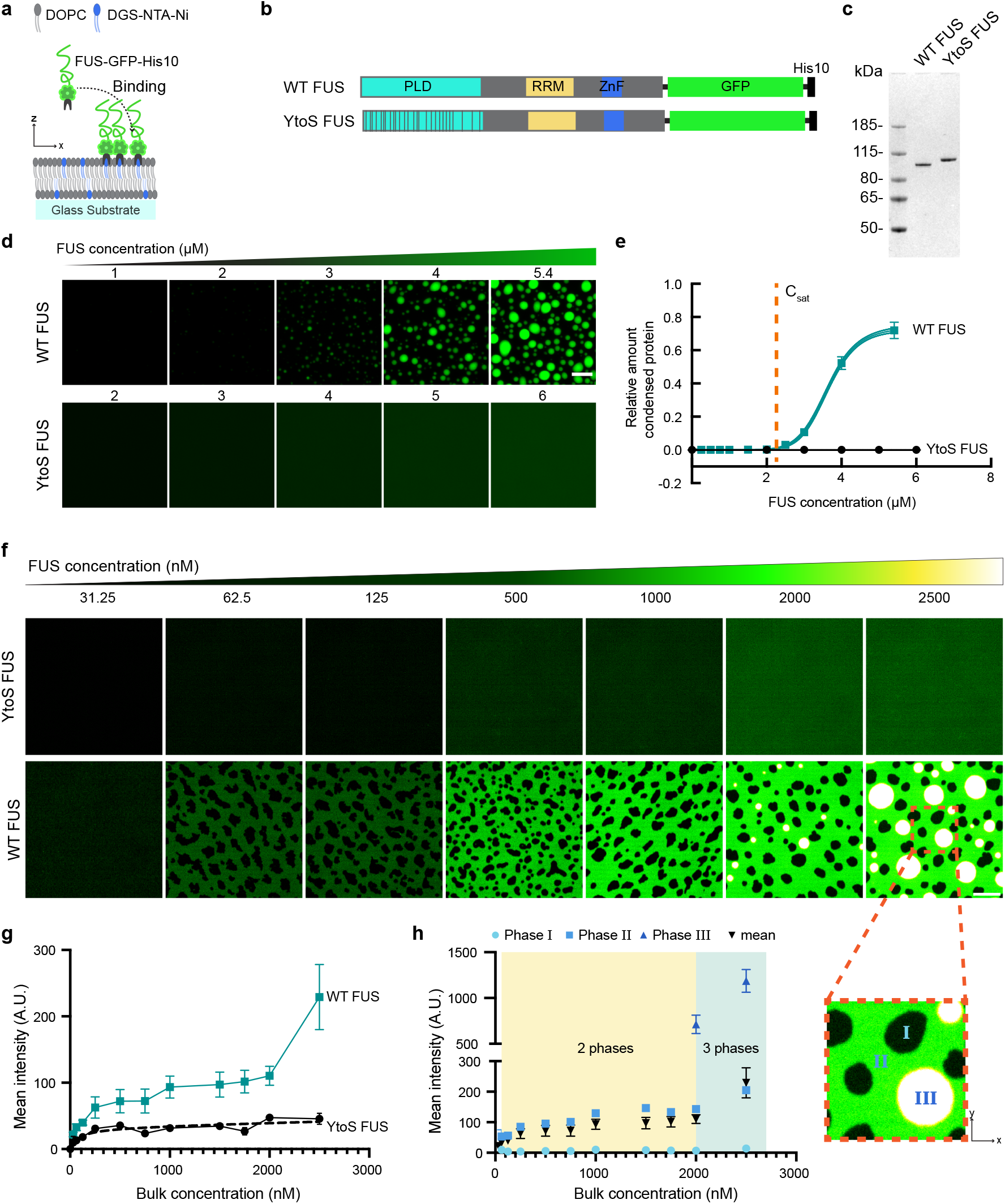
Binding to a lipid bilayer enables FUS to form 2D patterns at low concentrations. **a** Model of FUS-His10 binding to the lipid bilayer via His-Ni interactions. **b** Functional domains of the wild type FUS and its YtoS mutant, both with C-terminal GFP for visualization and a His10 tag at the C-terminus that enables binding to the lipid bilayer via NTA-Ni lipids. **c** SDS-PAGE of the two purified proteins. **d** Representative maximum intensity projection images of solutions of FUS protein at increasing concentrations confirming that the phase separation propensities or lack thereof for both wild type and YtoS mutant is not altered by additional tags. **e** Quantification of condensed protein for each construct relative to concentration. WT-FUS demonstrates a bulk saturation concentration *c*_sat_ = 2*µ*M, 3x independent experiments error bars = ± SD *n* = 9. **F** Representative images in the bilayer plane reveal that, at concentrations significantly lower than the bulk saturation concentration *c*_sat_, FUS can form 2D patterns. When the concentration reaches *c*_sat_, three phases coexist at the membrane (see the inset). **g** Binding curves showing the total amount of WT and YtoS protein accumulated at the bilayer before and after *c*_sat_ is achieved error bars = *±*SD *n* = 9. The dashed black line is the fitted dilute binding isotherm (Eq. (1)). **h** FUS intensity in the different phases error bars = *±*SD *n* = 9 3-phase coexistence of phases i ii and iii is exemplified in the inset image to the right.

In contrast, the YtoS construct did not form condensates under any of the tested concentrations up to 6µM. The two proteins were introduced to the supported lipid bilayers (SLB’s) composed of the lipid DOPC doped with 5 mol% of the receptor-like lipid DGS-NTA(Ni) (Figure 2a), which binds to the His10 tag of FUS with high affinity and specificity. In the absence of DGS-NTA(Ni), FUS was not observed to bind to the membrane, which allowed us to control membrane binding of FUS via the concentration of DGS-NTA lipid in the membrane. The non-phase separating YtoS mutant bound to the 5% DGS-NTA(Ni) supported lipid bilayers uniformly in space (Figure 2f, first row) and exhibited concentration-dependent Langmuir-like binding behavior, reaching a saturation point at which no more protein could bind (Figure 2g, black curve). The dashed black line in Figure 2g shows a fit to the concentration bound to the membrane surface *c*_*S*_ as a function of the bulk protein concentration *c*_*B*_ using the following dilute Langmuir isotherm

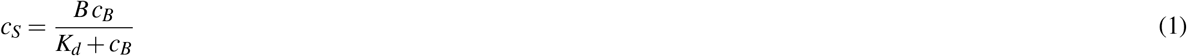

from which we extract the binding affinity *K*_*d*_ = (82.00 ± 36.10) nM. In contrast, membrane binding of WT-FUS could not be described with a classical binding-isotherm. At low bulk concentration the membrane bound density of WT-FUS increased beyond the levels of YtoS. At intermediate concentrations, WT-FUS membrane density did not saturate. Instead, the surface concentration appeared to increase linearly with a low slope until bulk *c*_sat_ was reached and the surface density drastically increased (Figure 2g, teal curve). This deviation from the YtoS behaviour also corresponds with the membrane becoming heterogeneous, reflected in two-dimensional FUS-WT patterns on the SLB’s (Figure 2f, second row). These two-dimensional patterns occurred for concentrations above 62.5 nM (what we termed the threshold concentration *c*_2D_), which was significantly lower than the experimental bulk saturation concentration *c*_sat_.

Above *c*_sat_, FUS-WT-rich droplets formed in the bulk and wetted the SLB’s. Interestingly, these wetted droplets coexist with the two-dimensional patterns corresponding to three-phase coexistence at the lipid bilayer, see Figure 2f (inset) and g. In Figure 2h, we quantify protein concentrations in the coexisting phases as a function of bulk protein concentration (see the Methods section for details on the quantification). In conclusion, despite having similar binding affinities for the membrane, the WT-FUS accumulated at a significantly higher concentration on the supported lipid bilayer than the YtoS mutant. Furthermore, WT-FUS showed 2D and 3D phase separation. The increase of the 2D dense phase concentration suggest that additional FUS is recruited from the bulk even after all membrane binding sites are saturated.

### FUS dense 2D phase recruits non-membrane-binding FUS

To test the hypothesis that the non-saturation behavior of WT-FUS at the membrane surface is related to recruitment of FUS from the bulk via FUS-FUS interactions instead of FUS-membrane interactions, an additional species of FUS-WT was generated that could not bind to the membrane and fluoresced at a different wavelength. A non-membrane-binding FUS was engineered by removing the His10 tag and adding a far-red fluorophore (Dylight 650) via amine-reactive labeling, thus minimally altering the size of the protein (Figure 3a). We confirmed that removing the His-tag significantly reduced the amount of protein recruited to the DGS-NTA bilayer alone (Figure 3b). The addition of the non-membrane binding FUS-WT-GFP-DL650 resulted in significant recruitment to the membrane when FUS-WT-GFP-His10 was bound to the bilayer(Figure 3b,d). Increasing the concentration of the non-binding species in the bulk caused a linear recruitment by the FUS bound to the bilayer. We speculate that the non-membrane binding proteins were recruited, leading to a multilayer of proteins. Moreover, this multilayer recruitment of proteins is driven by the interactions that drive phase separation of the protein.

**Figure 3.**
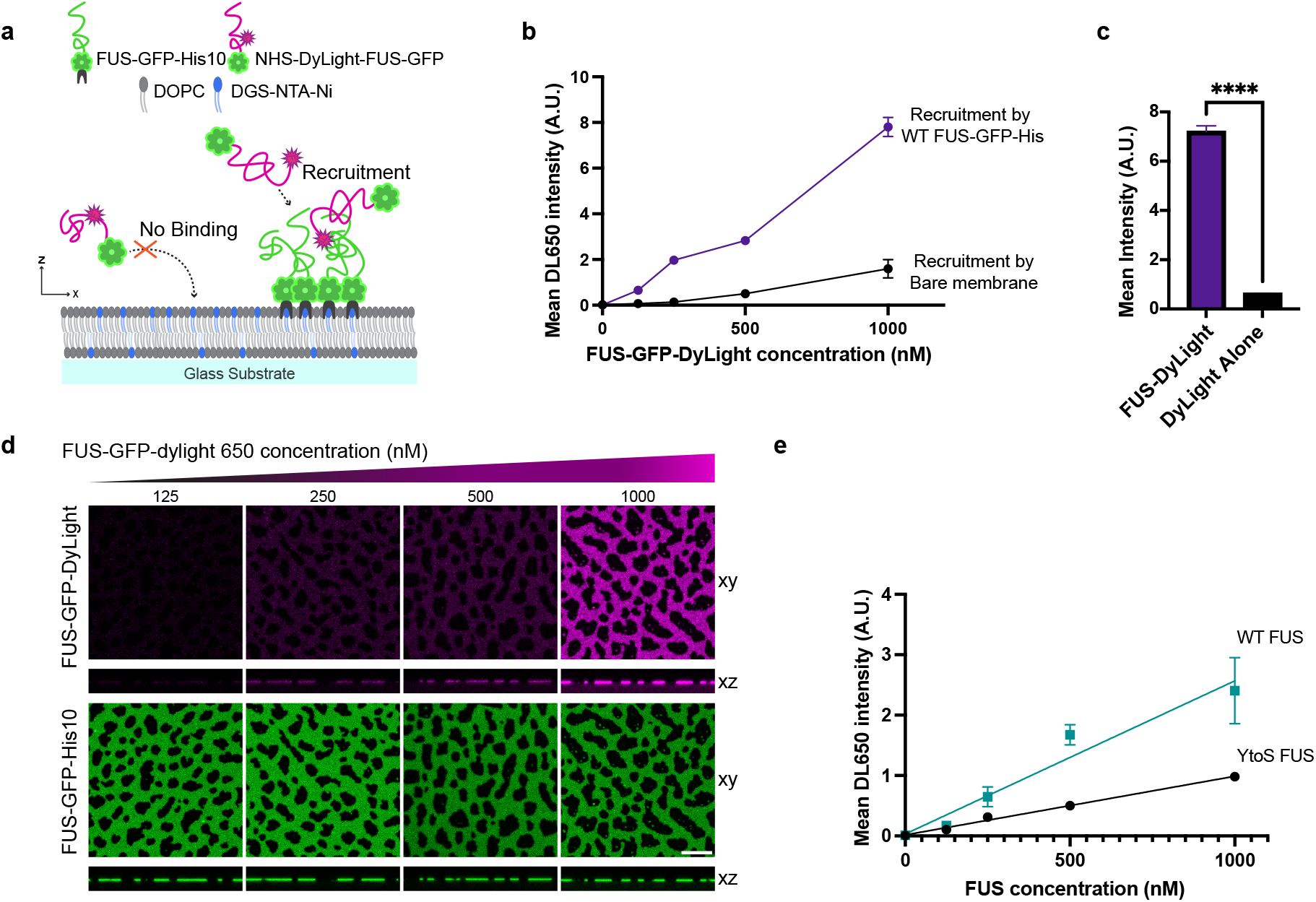
Two color assay suggests a multi-layer structure of the phase-separating FUS variant. **a** Model of FUS bound to the lipid bilayer recruiting WT-FUS-DL from solution via PLD-PLD interactions to form a multilayer of protein below bulk saturation concentration *c*_sat_. **b** Quantification of the WT-FUS-DL recruitment by bound WT FUS and the membrane alone, error bars = ± SD *n* = 4 measurements. **c** Quantification of 100nM WT-FUS-Dylight recruitment to membrane-bound FUS in comparison to an equivalent concentration of Dylight alone error bars = ± SD *n* = 4 measurements, unpaired t-test **** P<0.0001. **d** Representative confocal images in XY anx XZ plane of membrane-bound WT-FUS (green) recruiting non-binding WT-FUS labeled with dylight 650 (WT-FUS-DL magenta), scale bar 10 µm. **e** Comparison of non-membrane binding YtoS-DL and WT-DL to their respective binding equivalents, error bars = *±*SD *N* = 3 measurements.

To test this mechanism, we performed the same assay with the YtoS mutant and found that its ability to recruit more protein was significantly reduced with respect to the WT (Figure 3e). In order to rule out the effect of the additional fluorophore, we also tested the recruitment of the fluorophore alone at comparable concentrations. We found that its ability to be recruited was significantly less than the fluorophore with the protein (Figure 3c), confirming that the main driver of recruitment was the protein-protein interaction and not confounding effects of the labeling. Taken together, the results of this assay indicate the emergence of a multi-layer structure. This layer forms in two steps: First, high-affinity binding of the HIS10 tag to the NTA-lipid and second, recruitment of additional FUS molecules from solution via the FUS-FUS interactions. We hypothesize that the second step is mediated by the multivalent interactions via the tyrosines in the prion-like domain. Indeed, these interactions have been shown to drive phase separation in solution and lead to a non-stoichiometric binding behavior on the membrane^31^.

### 2D phase transitions are dependent on interaction strength

Having established that membrane binding of FUS leads to a surface phase transition with phase coexistence of dense and dilute domains on the membrane, we next aimed to assess whether this transition could be modulated in the same fashion as classical 3D bulk transitions, e.g. through changing the protein-protein interaction strength. A common method of modulating interaction strength is through varying the salt concentration^43–45^. To evaluate the effects of salt concentration on the 2D phase separation compared to 3D phase separation in bulk, we titrated the WT FUS protein on 5% Ni bilayers at salt concentrations ranging from 150mM KCl to 500mM KCl. As shown above, at 150mM KCl, a 2D phase separation emerged at FUS concentrations as low as 62.5nM, increasing the salt concentration and thereby lowering the interaction strength of the protein also increased the concentration *c*_2D_ at which 2D patterns emerge on the bilayer (Figure 4a). Further analysis at salt concentrations of 200mM, 300mM, and 500mM demonstrated that the recruitment of WT protein to the bilayer was significantly higher than that of YtoS at all protein concentrations (Figure 4b). Furthermore, the protein density in the dense phase increased with increasing bulk concentration (Figure 4b), in contrast to the behavior of simple binary mixtures.

**Figure 4.**
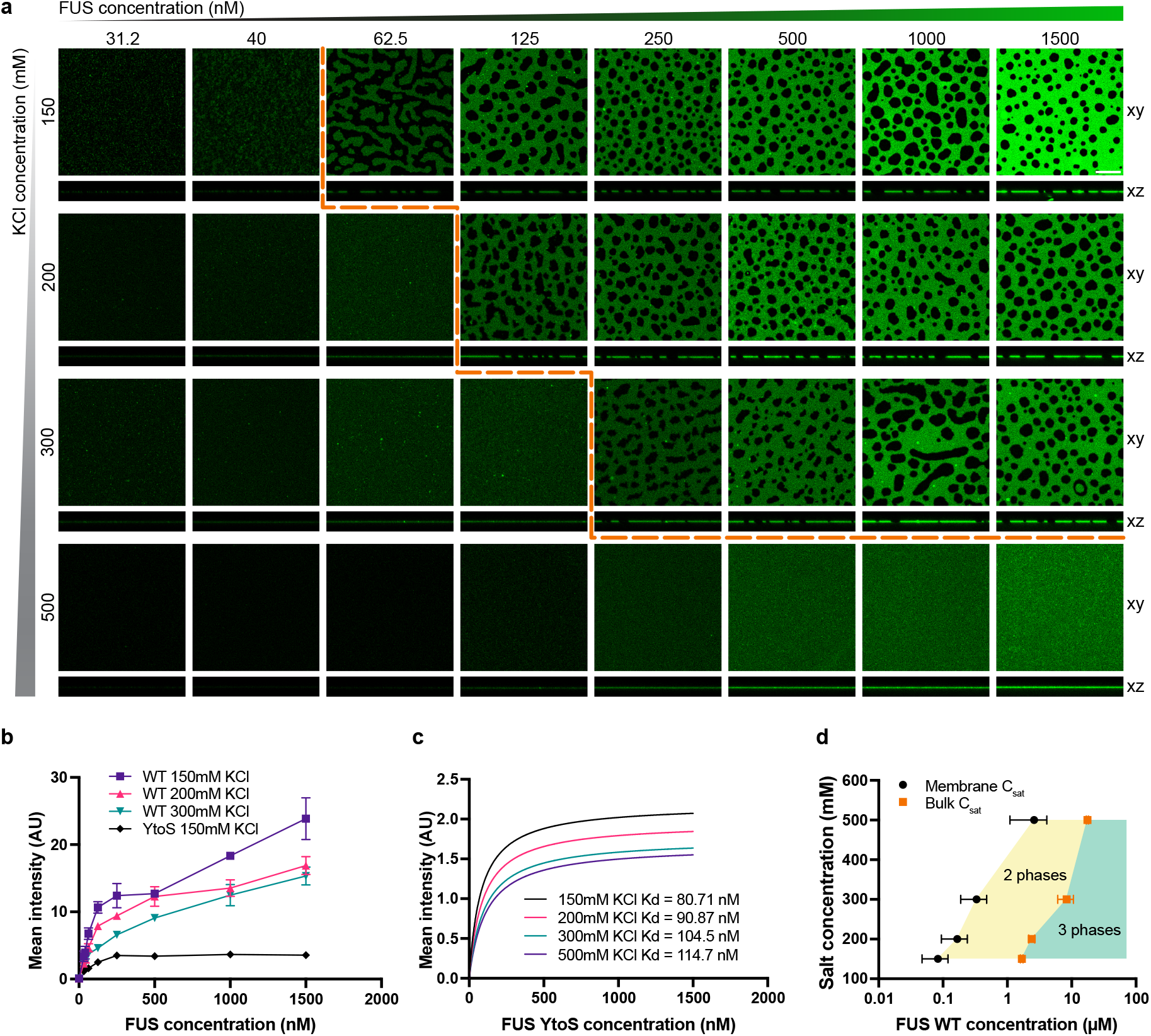
Phase diagram of the bulk-membrane coupled system. **a** Representative confocal images, with XZ profile beneath, of FUS titration on 5% Ni SLB’s in different salt buffer concentrations, demonstrating a shift in the onset of 2D phase separation with increasing salt, scale bar 10 µm. **b** Comparison of mean intensity of the dense phase of membrane-bound FUS under different salt concentrations with mean YtoS intensities for reference error bars = ± SD *n* = 4 measurements. **c** Comparison of apparent binding curves of YtoS under 150, 200, 300, and 500 mM KCl. **d** Comparison of the bulk *c*_sat_ (pink) with *c*_2D_, the concentration at which 2D patterns emerge (black), at differing salt concentrations. Error bars = ± SD *N* = 3 measurements.

To scrutinize whether salt concentration also affected the Ni-his tag interaction, binding curves for the lipid surface were calculated for the YtoS construct under the four salt concentrations (Figure 4c). Notably, increasing salt concentration does have a slight effect on the binding of the protein to the bilayer, in that the *B* (see Eq. Eq. (1)) decreases with increasing salt. Due to the nature of the experimental setup, the actual binding affinity could not be calculated as the Ni concentration was significantly higher than the actual *K*_*d*_. Therefore, the measured and calculated values along with that of Figure 2g coincide with the ‘Titration regime’^46^. Through increasing the salt concentration and, thereby modulating the binding and interaction strengths of the protein, we were able to tune *c*_2D_ and *c*_sat_, above which surface and bulk phase separation occur, respectively (Figure 4d). In particular, decreasing salt causes both *c*_2D_ and *c*_sat_ to decrease, thus making 2D and bulk patterns emerge at lower concentrations.

### A minimal model recapitulates the protein phase behaviour on membranes

To gain insights into how bulk protein and salt concentrations affect the emergence of coexisting phases on a surface, we constructed a thermodynamic model. The model considers a surface *S*, representing the lipid bilayer with an adjacent fluid layer of height *h* (Figure 5a). In the bulk, only two species are present: a bulk protein *B* and solvent *W*. In contrast, at the surface, three species are present: lipids without a receptor *L*, lipids tagged with a receptor *R*, and surface proteins *S*, i.e., proteins bound to lipid-receptor constructs. We make use of volume fractions for bulk species and area fractions for components bound to the surface. Specifically, we introduce the protein and solvent volume fractions, *φ*_*P*_ and *φ*_*W*_, respectively. The area fraction of proteins bound to receptor-lipid constructs is indicated with *ϕ*_*S*_, the area fraction of lipid-receptor constructs without proteins bound *ϕ*_*R*_, and the area fraction of untagged lipids with *ϕ*_*L*_. The phase behavior of the system is encoded in the bulk and surface free energy densities. In this work, we have used the following free energy density for the bulk species^47,48^:

**Figure 5.**
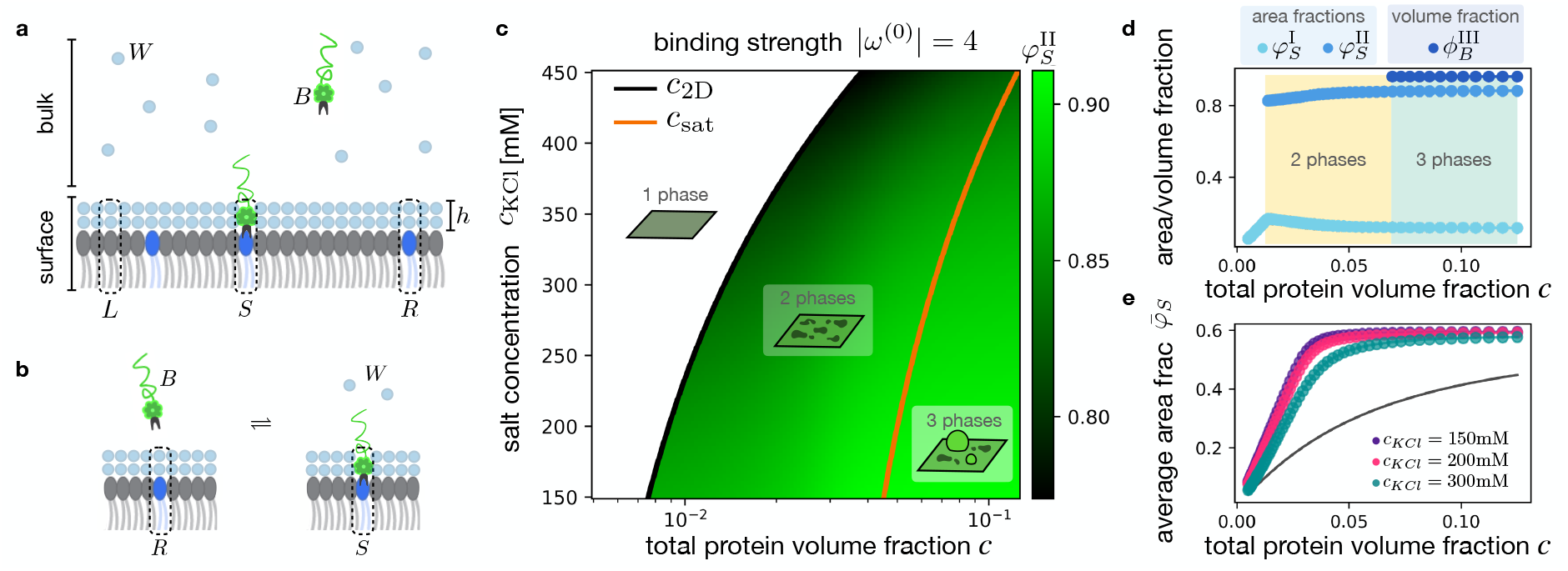
Theory of phase separation coupled to membrane binding recapitulates the FUS phase behaviour. **a** The model encompasses components located in the bulk and on the surface. The bulk components are protein (*B*), and solvent (*W*). The surface ones encompass lipids (*L*), and lipids tagged with a receptor (*R*), both including a solvent layer adjacent to the membrane, and protein bound to the surface via a lipid’s receptor (*S*). **b** Binding to the surface can be parametrized via a chemical reaction converting a bulk protein (*B*) and a tagged receptor (*R*) to a receptor membrane complex (*S*) while solvent (*W*) is released in the bulk. **c** Protein phase behaviour as a function of total protein volume fraction *c* and salt concentration, *c*_KCL_. The black and red curves indicate *c*_2D_ and *c*_sat_, i.e., the total protein volume fractions *c* at which phase separation occurs in the surface and bulk, respectively. The color code represents the area fraction of the membrane phase with the highest protein content, which we label as II. In qualitative agreement with the experimental phase diagram in Figure 4c, decreasing salt and increasing the protein amount favors coexisting phases on the surface. Increasing the protein amount leads to phase separation in the bulk as well. **d** Area fractions in the coexisting phases at the membrane, 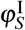and 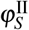, and in the bulk protein-rich phase 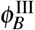 wetting the membrane, as a function of the total protein volume fraction. Here, *c*_KCL_ = 300 Nm. **E** Binding curve describing the area fraction of protein bound to the surface as a function of the total protein volume fraction *c*, for three representative values of salt. In black, we display the curve corresponding to a non-interacting protein, such as the YtoS mutant, i.e., we set *χ* = 0 and *χ*_*i j*_ = 0. For parameters, see the Methods section.

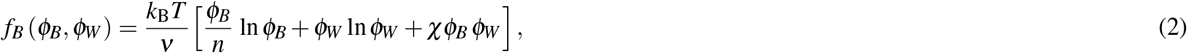

where *k*_*B*_*T* is the thermal energy, *ν* is a volume scale, *n* is the molecular volume ratio between proteins and solvent, and the interaction propensity χ quantifies protein-solvent interactions in the bulk. The free energy for the surface reads

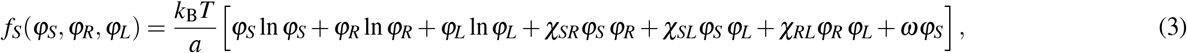

where the interaction propensities *χ*_*SR*_, *χ*_*SL*_, and *χ*_*RL*_ parameterize interactions between surface-bound proteins and free receptors, surface-bound proteins and untagged lipids, and free receptors and untagged lipids, respectively. Furthermore, we have introduced *ω*, the so-called internal free energy, quantifying the free energy released upon a binding event in the dilute limit, and the area scale *a* = *nν/h*.

The interaction propensities and internal free energy are affected by various physical control parameters, including temperature, pH, and salt^13,49^. In our work, we focus on the effect of the concentration of the KCL salt, *c*_KCL_. To this end, we write a phenomenological and linear relationship of the interaction parameters for bulk and membrane surface, and the internal free energy:

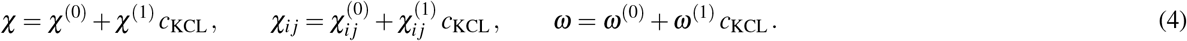

Specifically, we consider the case where increasing the salt concentration decreases the ability to phase separate in the bulk (*χ*^(1)^*c*_KCL_ *<* 0) and in the membrane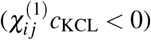, and lowers the affinity for binding (*ω* less negative by choosing *ω*^(1)^ *c*_KCL_ *>* 0).

Total material conservation in the bulk and at the surface implies that

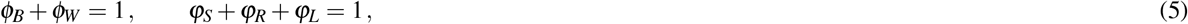

reducing the amount of independent volume and area fractions from five to three, which we chose to be *φ*_*B*_, *ϕ*_*S*_, and *ϕ*_*R*_. Molecular binding can be represented as a chemical reaction among the components (Figure 5b):

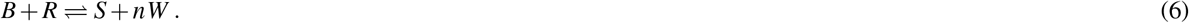

At binding equilibrium, the number of independent fields is further reduced to the two quantities conserved by the reaction in Eq. (6), namely the total protein volume fraction *c* = *φ*_*B*_ + *ϕ*_*S*_ *hA/V* and total amount of untagged lipid *ϕ*_*L*_^35^. Here, *h* and *A* are the bilayer height and area, respectively, and *V* is the system volume. Both conserved quantities of total protein volume fraction *c* and lipid amount without receptor *ϕ*_*L*_ are accessible in the experiments. Indeed, *c* corresponds to the volume fraction of protein used in the preparation of the experimental system. After preparation, a fraction of the total protein amount binds to the membrane surface *ϕ*_*S*_, while the remaining amount *φ*_*B*_ = *c* − *ϕ*_*S*_ *hA/V* remains in the bulk. Moreover, the area fraction of lipids without receptors *ϕ*_*L*_ can be easily controlled during the preparation of the lipid bilayer by varying the amount of DGS-N/TA(Ni) lipid. As explained in detail in the Methods section, our model enables us to determine the average protein volume fraction in the bulk *φ*_*B*_ and average area fractions of the receptor-protein constructs at the surface *ϕ*_*S*_, a function of the conserved quantities *c* and *ϕ*_*L*_. Moreover, if phase separation at the surfaces and/or in the bulk occur, we get the area fraction in the two phases coexisting at the surface, i.e., 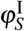 and 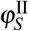, as well as the volume fractions of the bulk phases, i.e., 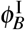 and 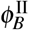. For all subsequent studies, we fix the amount of lipids without receptor *ϕ*_*L*_ and consider the conserved total protein volume fraction *c* and the salt concentration *c*_*KCl*_ as the system’s control parameters.

In Figure 5c, we display the phase diagram as a function of total protein volume fraction *c* and salt concentration *c*_*KCl*_ for a fixed lipid composition *ϕ*_*L*_. The salt concentration affects the interaction and binding strength (see Eq. (4)). Specifically, the propensity for phase separation in bulk and membrane surface decreases, and the binding affinity reduces with increasing salt concentration. The black and orange lines represent *c*_2D_ and *c*_sat_, i.e., the total protein volume fraction at which phase separation occurs at the surface and in the bulk, respectively. In the white domain, both the bulk and the membrane are homogeneous. Between the black and the orange lines, proteins bound to the surfaces phase separate. For concentrations larger than *c*_sat_, bulk proteins can phase separate as well, and, assuming partial wetting, three phases can coexist at the membrane. The color code indicates the area fraction 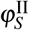 in the protein-rich phase at the membrane.

We summarize that the model leads to a phase diagram that is consistent with experiments (compare to Figure 5c and Figure 4d). In Figure 5d, we show the protein area fraction in coexisting phases at the membrane, 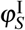 and 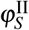, and the protein volume fraction in the protein-rich bulk phase 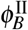, as a function of the total protein volume fraction *c*. Here, *c* _*KCL*_ = 150*m*M. Note that the curves in Figure 5d are in good qualitative agreement with the experimental results in Figure 2h. Lastly, in Figure 5e, we show how salt concentration influences the average area fraction of protein bound to the surface *ϕ*_*S*_, as a function of the total protein volume fraction *c*. In agreement with the experimental results in Fig. 4b, increasing salt leads to a decrease in the amount of protein bound to the surface.

To understand how the gap between the 2D and 3D binodal (i.e., the region in the phase diagram where surface but no bulk phase separation occurs) is affected by the affinity of binding to the membrane surface, we quantified how varying binding strength |*ω*^(0)^ |alters the onset of two and three-phase coexistence. Strictly speaking, *ω*^(0)^ is the internal energy in the absence of salt, but its absolute value |*ω*^(0)^ |quantifies the binding and is therefore referred to as binding strength. The larger the binding strength, the more proteins are bound. In Fig. 6a-c, we show two phase diagrams corresponding to relatively low and high binding strengths, respectively. At lower binding strength, the onset of 2D phase separation shifts to high values of the total protein volume fraction *c* (Fig. 5a). At high salt concentration, bulk phase separation occurs before membrane phase separation, i.e. *c*_sat_ *< c*_2D_. For concentrations *c* such that *c*_sat_ *< c < c*_2D_ bulk droplets wet the surface, but no 2D phases are found. As a consequence, the three-phase coexistence region shrinks. At high binding strength (Fig. 5b), the onset of 2D phase separation shifts to lower values of the total protein volume fraction *c*, widening the region where 2D patterns are found at the membrane but not in the bulk. When increasing the binding strength |*ω*^(0)^ | (from Fig. 5a to b), most receptor are bound. As a result, salt has fewer effects on *c*_2D_, corresponding to the black line getting more vertical.

**Figure 6.**
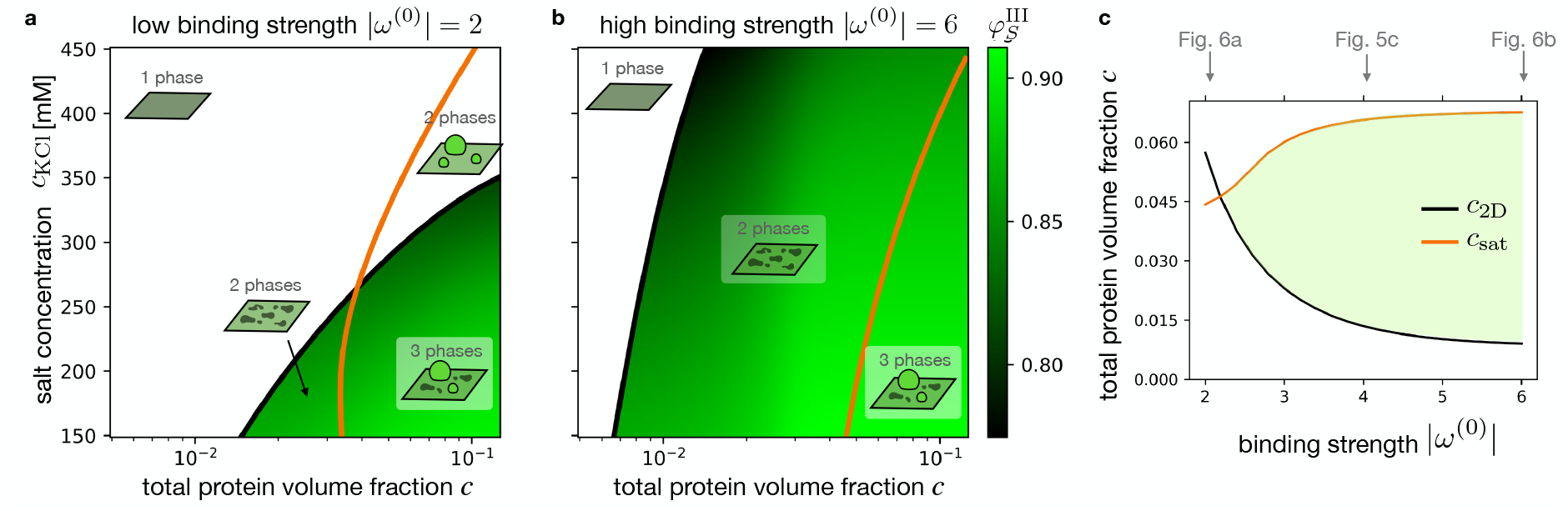
Binding affinity tunes the onset of two- and three-phase coexistence. Theoretical phase diagrams for low **a** and high **b** binding strength *ω*^(0)^, as a function of total protein volume fraction *c* and salt *c*_KCL_. As before, we indicate *c*_2D_ (black curve) and *c*_sat_ (orange curve), which are the total protein volume fractions *c* at which 2D and bulk phase separation occur. Interestingly, for low binding strength |*ω*^(0)^ |, high salt concentrations lead to bulk in the absence of surface phase separation, with *c*_sat_ *< c*_2D_. **c** *c*_2D_ and *c*_sat_ as a function of the binding strength |*ω*^(0)^ |, for a fixed salt concentration *c*_KCl_ = 300 nM. For large binding strength |*ω*^(0)^ |, *c*_2D_ decreases to a constant and non-zero value that is set by the total receptor concentration since all receptors are bound. Moreover, for large |*ω*^(0)^ |, the bulk saturation concentration *c*_sat_ slightly shifts to larger total protein volume fractions *c*, since more molecules are bound, which slightly depletes the molecules in the bulk. A much more pronounced effect is observed for *c*_2D_ when increasing |*ω*^(0)^ | since more recruitment of molecules to the membrane favors phase separation in the membrane.

In Fig. 5c, we fix the salt concentration to *c*_KCL_ = 300*m*M and quantify the effect of varying binding strength on the total concentration *c* at which 2D and 3D phase separation occurs, i.e., *c*_2D_ and *c*_sat_, respectively. For our parameter set, at low binding strength, *c*_sat_ *< c*_2D_. Increasing binding strength leads to a decrease of *c*_2D_ (Fig. 5c, dashed line), while *c*_sat_ remains almost constant. This behavior leads to a pronounced reduction of *c*_2D_ to very low protein volume fractions. This trend effectively enables 2D phase separation at the membrane surface over a wide range of protein volume fractions, preceding the onset of 3D phase separation by a considerable margin. We conclude that the membrane binding affinity is a sensitive parameter that can enable surface phase separation at concentrations orders of magnitude below 3D bulk phase separation.

## Discussion

Here we used a minimal model system to study protein phase transitions in the context of membrane binding. Using the well-characterized phase-separating protein FUS as our model protein and modifying it to have the capability of specific binding to a lipid bilayer membrane, we were able to explore the onset of 2D surface phase separation at subsaturated concentrations (i.e., concentrations at which no bulk condensates are present). Our main findings are that high-affinity membrane binding of a cytosolic multivalent protein leads to the emergence of 2D phase separation that occurs at concentrations which are an order of magnitude lower than the bulk saturation concentration *c*_sat_. For concentrations above *c*_sat_, 3-phase coexistence can occur at the lipid bilayer, which has been observed in many other systems before^14,33,39,50–58^. In addition, our results show that as bulk protein concentration approaches *c*_sat_, the 2D dense phase increases by recruiting additional protein from the bulk via protein-protein interactions. This trend suggests the formation of a thicker multilayer of protein emerges on the lipid bilayer membrane close to the 3D phase boundary. Increasing the bulk concentration results in an increase in phase concentrations and volume fraction, in contrast with binary phase separation in the bulk, where only the phase fractions change.

Furthermore, using thermodynamic theory, we quantified the concentrations at which 2D and 3D phase separation occur (*c*_sat_ and *c*_2D_) as a function of essential thermodynamic control parameters, such as salt concentration and binding strength. Our results show that the range in protein concentration leading to surface phase separation in the phase diagram increases and shifts to very low protein concentration by increasing the binding affinity to the membrane. This mechanism can provide a robust switch to control phase separation in cells by tuning the binding affinity of cytosolic scaffold proteins to activated receptors or signaling lipids. Along these lines, cells can express scaffold proteins at concentrations far below the bulk saturation concentration but still turn on phase separation locally at the membrane by enabling context-dependent high-affinity binding to the membrane, for example, via receptor oligomerization, phosphorylation, calcium secretion, or other signaling-related membrane changes. In this scenario, spontaneous (uncontrolled) phase separation in the bulk is extremely unlikely even in a fluctuating environment such as the cytoplasm. At the same time, small signaling events at the membrane can be robustly translated into a phase transition to build membrane-attached compartments that amplify signaling or drive assembly of super-molecular membrane structures. This scenario can be applied to many biological processes, in which phase separation at the membrane has been shown to play a role (endocytosis^52^, T-cell activation^14,53^, post-synaptic density^54,55^, tight junction formation^33,56–58^). However, so far, the role of membrane binding affinity in the phase transition has not been studied in isolation, because in most of these systems, the interactions (multivalence) of the scaffold protein and the membrane binding affinity are altered to achieve the phase transition. By combining the simplicity of our model system with theory, we demonstrate that tuning the membrane binding affinity is sufficient to enable robust local phase separation.

## Methods

### Protein expression and purification

Plasmids containing WT and YtoS FUS constructs^42^ were generously provided by the Hyman lab. New constructs were generated by restriction cloning the WT and YtoS FUS into the FlexiBAC shuttle vector pOCC143-pOEM1-N-MBP-3C-C-mGFP-TEV-HIS10 using NotI/AscI restriction sites and Quick Ligase™. Baculovirus for the expression of constructs was generated by the MPI-CBG protein expression and chromatography facility using methods previously described^59^. SF+ cells were infected with 2% by volume of baculovirus and incubated at 26.5C for 48 hours. Cells were collected through centrifugation and lysed using an LM20 Microfluidizer at 12000psi. The proteins were affinity purified sequentially via IMAC using Qiagen Ni-NTA agarose resin then NEB amylose resin. The MBP tag was removed using in-house 3C-protease digest and the His10 tag, when desired, was removed using NEB TEV protease. The desired protein was then purified from the cleaved tags via SEC on an Äkta Pure protein purification system with a HiLoad 16/600 Superdex200 pg column.

The purification buffers were as follows: Main Buffer-1M KCl, 20mM HEPES, pH 7.25. Lysis Buffer-Main Buffer supplemented with 20mM Imidazol, 1mM MgCl_2_, Roche EDTA free protease inhibitor cocktail. His Wash Buffer-Main Buffer supplemented with 20mM Imidazol. His Elution Buffer-Main Buffer supplemented with 500mM Imidazol. MBP Wash Buffer-Main Buffer supplemented with 5% glycerol and 1mM DTT. MBP Elution Buffer-MBP Wash Buffer supplemented with 20mM Maltose. SEC Buffer-0.5M KCl, 20mM HEPES, pH 7.25, 5% glycerol, 1mM DTT.

### Supported Lipid Bilayer generation

SLB’s were generated using small unilamellar vesicles(SUV’s). In short, SUV’s were generated by preparing a lipid mixture (0.02mol% fluorescent lipid (DPPE-KK114(abberior STRED-0200-500UG) or 18:1 Lis-Rhodamine PE(Avanti 810150)), 0.1% 18:1 PEG5000 PE(Avanti 880230), 5% 18:1 DGS-NTA(Ni)(Avanti 790404), 94.88% DOPC(Avanti 850375)) in chloroform, drying the mixture overnight under vacuum, resuspending the lipids to a total concentration of 1mM in SUV buffer (150mM KCl 20mM HEPES pH 7.25), freeze thawing the resuspended lipids 10x by cycling between liquid N_2_ and a 42C water bath, and extruding them using an Avanti mini-extruder kit(Avanti 610000) and polycarbonate membrane with 100nm pore size.

A Greiner 96-well glass bottom cellview microplate(655891) was prepared by incubating in 2% Hellmanex III(Hellma 9-307-011-4-507) overnight, then 2×30-minute incubations in 5M KOH at 42C the following day. After the plate was extensively washed in distilled water and dried, the SUV’s were diluted to a final concentration of 300µM in 50µL for each well and incubated at room temperature for 1 hour. 30µL of 4M KCl was subsequently added to each well and the plate was washed with SLB buffer (11.2mM KCl 20mM HEPES pH 7.25) using a Tecan HydroSpeed™ plate washer with a custom programme designed to leave a final volume of 50µL buffer in each well. The quality of each bilayer was assessed through FRAP. Recovery of fluorescence in a bleach spot as well as a homogeneously fluorescent surface were indicative of a high quality bilayer.

### Bulk phase separation assay

Protein was diluted to various concentration in the desired salt buffer to a total solution volume of 25µL. The sample was then added to an ultra-low-binding 384-well plate(revvity 6057800). Condensates were left to settle at the bottom of the plate for 2 hours. For the comparison of Bulk phase separation of WT vs YtoS FUS at 150mM KCl, images for each well were acquired using an Olympus IXplore Spin SR SD, high resol, inverted IX83, Yokogawa W1 CSU with a 40x 0.95NA UplanS Apochromat air objective. For the assessment of saturation concetration at different salt concentrations, images for each well were acquired using an Olympus FLUOVIEW 3000, inverted IX83 stand with a 20x 0.8NA U Plan X-Apochromat air objective. Sum projections were generated and the relative amount of condensed protein vs protein concnetration was determined using the previously established custom built python analysis pipeline^42^.

### High throughput membrane binding and recruitment assay

A custom high throughput assay was developed in collaboration with the MPI-CBG Technology development studio (TDS). Protein was asdded to each well of a greiner 96 well plate containining SLB’s and left for 1 hour to equilibrate at room temperature. The plate was then imaged using a Yokogawa CV7000 high-content spinning disc confocal microscope with a 60x 1.2NA water immersion objective. Images were acquired at the bilayer and 20µm above in solution. The resultant images were analyzed using a custom python script to differentiate between phases and quantify their intensities.

### 2-colour assay

Protein that had been digested with 3C and TEV protease was substoichiometrically labelled with DyLight 650 NHS Ester(Thermo 62265) through amine reactive labelling following the manufacturer’s protocol. Labeling reactinos were performed in a 1:2 dye:protein ration resulting in a labelling efficiency of 30-45%. For the assay, the labeled protein was diluted to a final labeling of 5% in the double digested GFP tagged protein. 5% Ni membranes were generated with Rhodamine-PE as the fluorescent lipid and FUSGFP-His10 (WT or YtoS)was added to the well to bind and saturate the membrane (concentrations>250nM) for 1 hour. The remaining protein was rinsed 5 times with 150mM KCl SLB buffer, the DyLight-FUS-GFP protein was titrated from 125 to 1000nM, and incubate at room temperature for 30 minutes before imaging. Images were acquired using a confocal infinty-line STED microscope (Abberior Instruments) operated using the Imspector software (v.16.2.8415), with pulsed laser excitation (490nm, 560nm, 640nm, 40MHz), and ×60 water(Olympus). Images were analyzed using a Fiji^60^ macro that created a mask (gaussian blur (sigma=1) and default thresholding where the black background was set to false) that was then used to measure the fluorescence intensity within the dense regions of protein as well as the total intensity of the entire image including dilute and dense phases.

### 2D saturation concentration dependence on salt

5%Ni SLB’s were prepared in a 96-well greiner plate and protein was titrated final concentrations between 31.2nM and 1500nM (final volume 70µL) in 150, 200, 300, and 500mM KCl. Protein was left to equilibrate for 1 hour at room temperature before imaging with the commercial confocal STED microscope described previously. In order to determine the 2D saturation concentration concentration at 500mM, protein was purified in 1M KCl and titrated to final concetrations between 54nM and 21.95µM. protein was left to equilibrate 1 hour and images were acquired using an Olympus FLUOVIEW 3000, inverted IX83 stand with a U Plan S-Apo 60x (1.2 NA) water objective. Images were visually assessed for the presence of 2 phases to check for the transition point from single to multiple phases and were quantified using Fiji^60^ and the macro described above.

### Phase and binding equilibria

We now introduce the bulk chemical potentials, *µ*_*B*_ = *nν* ∂ *f*_*B*_*/*∂*φ*_*B*_ and *µ*_*W*_ = *ν* ∂ *f*_*B*_*/*∂*φ*_*W*_, and the surface chemical potentials *µ*_*S*_ = *a*∂ *f*_*S*_*/*∂*ϕ*_*S*_ and *µ*_*R*_ = *a*∂ *f*_*S*_*/*∂*ϕ*_*R*_, and *µ*_*L*_ = *a*∂ *f*_*S*_*/*∂*ϕ*_*L*_. Imposing binding equilibrium, namely

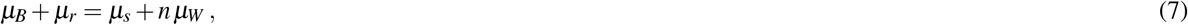

together with the conservation laws in Eq. (5), allows us to find the volume fractions of all the components, *φ*_*B*_, *φ*_*W*_, *ϕ*_*S*_, and *ϕ*_*R*_ as a function of the quantities conserved in the chemical reaction, namely *c* = *φ*_*B*_ + *ϕ*_*S*_ *hA/V* and *ϕ*_*L*_.

Then we can look for phase coexistence in the bulk and the surface. Defining Π_*S*_ = − *f*_*S*_ + *ϕ*_*S*_ ∂ *f*_*S*_*/*∂*ϕ*_*S*_ + ∂ *f*_*R*_*/*∂*ϕ*_*R*_. The conditions for two phases at the surface I and II to stably coexist, read^61^:

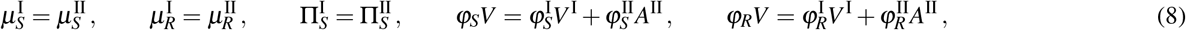

where *A*^I^ and *A*^II^ are the areas of phases I and II, respectively, that sum up to the total surface area *A* = *A*^I^ + *A*^II^. Defining now Π_*B*_ = −*f*_*B*_ + *φ*_*B*_ ∂ *f*_*B*_*/*∂*φ*_*B*_, the conditions for two bulk phases III and IV to stably coexist, read:

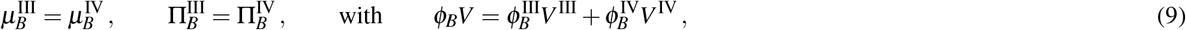

where *V* ^III^ and *V* ^IV^ are the volumes of phases I and II, respectively, that sum up to the total system volume *V* = *V* ^III^ +*V* ^IV^. In the main text, we have labelled I and II the protein-poor and protein-rich phases at the surface, respectively. We also labelled III the protein-rich bulk phase, which, in experiments, corresponds to droplets wetting the bilayer. Finally, note that, at binding equilibrium, the Gibbs phase rule imposes a maximum of 4 coexisting phases^62,63^. The parameters used in Fig. 5 and Fig. 6 are: *ϕ*_*L*_ = 0.4, *n* = 1, *χ*^(0)^ = 5, *χ*^(1)^ = −1*/*200 mM^−1^, 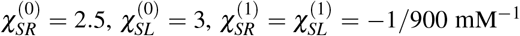, with the other interaction propensities are set to 0. We note that the interaction parameter in the bulk decreases (*χ*^(1)^ *<* 0), corresponding to the fact that increasing salt reduces the propensity to phase separate in 3D. Moreover, 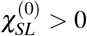, meaning that salt also lowers phase separation in the membrane. The internal free energy *ω*^(0)^ is indicated in the panels, while we set *ω*^(1)^ = 1*/*200 mM^−1^, corresponding to the situation where the binding propensity to the membrane reduces as the salt concentration increases. Furthermore *hA/V* = 0.05. In Fig. 5 d the salt concentration is 300 mM.

### Plotting, fitting, and statistics

GraphPad Prism and Python were used for plotting, fitting and performing statistical analysis on data.

## Acknowledgments

We thank Patrick McCall, Tyler Harmon, Frank Jülicher, and Stephan Grill for insightful discussions. G. Bartolucci acknowledges the Agencia Estatal de Investigación for funding through the Juan de la Cierva postdoctoral programme JDC2023-051554-I. X. Zhao acknowledges the “FoSE New Researchers Grant” of the University of Nottingham Ningbo China for financial support. We thank the PEPC facility and TDS at MPI-CBG for protein purification and high throughput imaging respectively. D. Sun acknowledges support by a seed grant of the DFG funded “Physics of Life” Excellent Cluster at TU-Dresden. A. Honigmann and C. Weber acknowledge the SPP 2191 “Molecular Mechanisms of Functional Phase Separation” of the German Science Foundation for financial support and for providing an excellent collaborative environment.

## Author contributions statement

I. L.-B. performed the experiments. G.B., X.Z., and C.A.W. developed the model. G.B. and I.L.-B. analyzed the data. All authors discussed and interpreted the results, and contributed to the writing of the manuscript.

